# The Influence of Ionic Environment on Nucleosome-Mica Interactions Revealed via Molecular Dynamics Simulations

**DOI:** 10.1101/2024.06.25.600666

**Authors:** Nilusha L. Kariyawasam, Jeff Wereszczynski

## Abstract

Mica serves as a crucial substrate in Atomic Force Microscopy (AFM) studies for visualizing and characterizing nucleosomes. Nucleosomes interact with the negatively charged mica surface via adsorbed cations. However, the specific influences of monovalent and divalent cations on nucleosome adsorption to the mica surface remain unclear. In this study, we investigated the binding of nucleosomes to the mica surface in the presence of monovalent potassium ions and divalent magnesium ions using molecular dynamics simulations. We also explored the impact of pre-treated mica surfaces on nucleosome binding and structure. Our findings reveal that nucleosome-mica interactions vary depending on the cations present, resulting in distinct effects on nucleosome structure. Notably, nucleosomes bind effectively to a mica surface in the presence of potassium ions with minimal structural perturbations.

## Introduction

The fundamental unit of chromatin is the nucleosome, which consists of eight core histone proteins and an approximately equal amount of DNA. The nucleosome core particle (NCP) is composed of a protein octamer made up of two copies each of the four core histones: H2A, H2B, H3, and H4.^1,2^ These histones physically associate into two H2A/H2B dimers and an (H3/H4)_2_ tetramer. Approximately 145-147 base pairs of DNA wrap around this complex. This nucleosome structure is essential for packaging and organizing DNA within the nucleus.^3^ Recently, it has been shown that nucleosomes are highly dynamic structures, undergoing processes over a vast range of lengths and timescales, allowing DNA and protein accessibility crucial for processes such as transcription, DNA replication, and repair.^4–6^

Atomic force microscopy (AFM) experiments can directly provide nanometer-scale descriptions of large-scale conformational changes in biomolecular complexes, and high-speed AFM (HS-AFM) can achieve this with millisecond temporal resolution.^7–9^ These techniques are highly effective for studying nucleosomes, as they enable visualization and characterization of both mononucleosomes and longer arrays.^10^ For example, HS-AFM has been used to visualize DNA sliding and unwrapping from nucleosomes at millisecond timescales.^7,8,11^

Both traditional and high-speed AFM require a flat surface that attracts biomolecules. Muscovite mica (KAl_2_(AlSi_3_)O_10_(OH)_2_), which becomes negatively charged in liquid, is a convenient substrate for immobilizing biomolecules via electrostatic interactions. ^12^ AFM studies have shown that the attraction between the mica surface and negatively charged DNA is mediated by cations present in the system.^13,14^ The mica surface can be pre-treated by replacing potassium ions with other cations such as Ni^2+^, Mg^2+^, Ca^2+^, Co^2+^, La^3+^, and Zr^4+^, which forms a more stable bridge between the negatively charged mica and DNA. ^13,15–17^ Additionally, the mica surface can be functionalized with 1-(3-aminopropyl) silatrane (APS) to create a weakly cationic surface.^18,19^ Several studies have aimed to understand how DNA binds to mica surfaces and how this process is modulated by solution cations. Hansma and coworkers demonstrated that DNA binding to a mica surface is correlated with the radius of transition metal cations in the buffer solution, based on their ability to fit into the cavities on the mica surface.^13^ Their study showed that DNA binds more effectively to mica when Ni^2+^, Co^2+^, and Zn^2+^ are present, which have ionic radii ranging from 0.69 to 0.74 Å. Despite having a similar ionic radius to Ni^2+^, Mg^2+^ does not bind DNA as strongly, potentially due to Mg^2+^ being a Group 2 metal ion with p electrons, while transition metals have higher enthalpies of hydration, which form stronger complexes.^13^ Pastre and coworkers have shown that DNA binding to mica can be enhanced when the mica is pre-treated with transition metal ions (Ni^2+^, Zn^2+^).^14^ Their study also indicated that monovalent cations are not effective in DNA attraction, and the high ionic strength of divalent cations weakens binding strength due to ion screening. Reversible DNA binding to mica is preferred to preserve native DNA-protein interactions while allowing imaging. AFM studies have shown that a combination of divalent and monovalent ions (Mg^2+^, K^+^) with pre-treated mica achieves reversible binding.^12,20^ In contrast, Ellis and coworkers found that monovalent ions (K^+^, Na^+^) in the buffer can attract DNA to the mica surface with minimal DNA-surface interactions and short immobilization times. They suggested that immobilization with monovalent cations may be beneficial because it allows conformational freedom of the biomolecule with weak surface interactions.^21^

Despite these studies, the molecular roles of divalent and monovalent cations on nucleosome adsorption to mica surfaces are not fully understood. In particular, it remains unclear how cations and mica surface binding alter the structure and dynamics of nucleosomal DNA and the histone tails. These changes are particularly important for the interpretation and analysis of AFM experiments, as molecular-level differences in the structure and dynamics of nucleosomes at these surfaces from solution states may not be apparent from the relatively coarse AFM images. To address this, we employed all-atom molecular dynamics simulations to investigate the process of nucleosome binding to the mica surface. Through our computational models, we explore the effects of different cations and the effects of regular or pre-treated mica surface on nucleosome adsorption, as well as the implications of the mica surface on the structure and dynamics of nucleosomal DNA and histone tails. Our simulation study indicates that nucleosome binding affinity to the mica surface varies under different ionic environments. Moreover, the structure and dynamics of the nucleosome can be affected by interactions with the mica surface, depending on the ionic environment. Our results suggest that nucleosomes in KCl solution at physiological concentration with a regular mica surface provides a suitable environment for nucleosome adsorption with minimal perturbation to the nucleosome structure.

## Methods

### DNA Metadynamics Simulations

A double-layer muscovite mica surface with dimensions of 72 Å *×* 72 Å *×* 20 Å was generated using CHARMM-GUI,^22^ and a B-DNA dodecamer, based on PDB 1JGR,^23^ was placed 20 Å above it. The initial mica structure contained potassium ions evenly distributed on both sides of the surface. An 80 Å-thick water box was constructed on top of the mica surface using Visual Molecular Dynamics (VMD).^24^ NaCl, KCl, and MgCl_2_ salts were used to neutralize and set the salt concentration to 150 mM. A magnesium pre-treated mica surface, denoted as mica ptd, was obtained by equilibrating a mica surface in MgCl_2_ solution for 4 µs. The CHARMM36 force field^25^ was used for the DNA, the mica surface was parameterized via the INTERFACE force field, ^26^ and the ions were treated with CUFIX parameters developed for the CHARMM force field^27^ with the TIP3P water model.^28,29^ Hexa-hydrated magnesium ions ([Mg(H_2_O)_6_]^2+^) were used.^30^ A summary of the systems prepared for metadynamics simulations is presented in Table 1.

**Table 1:**
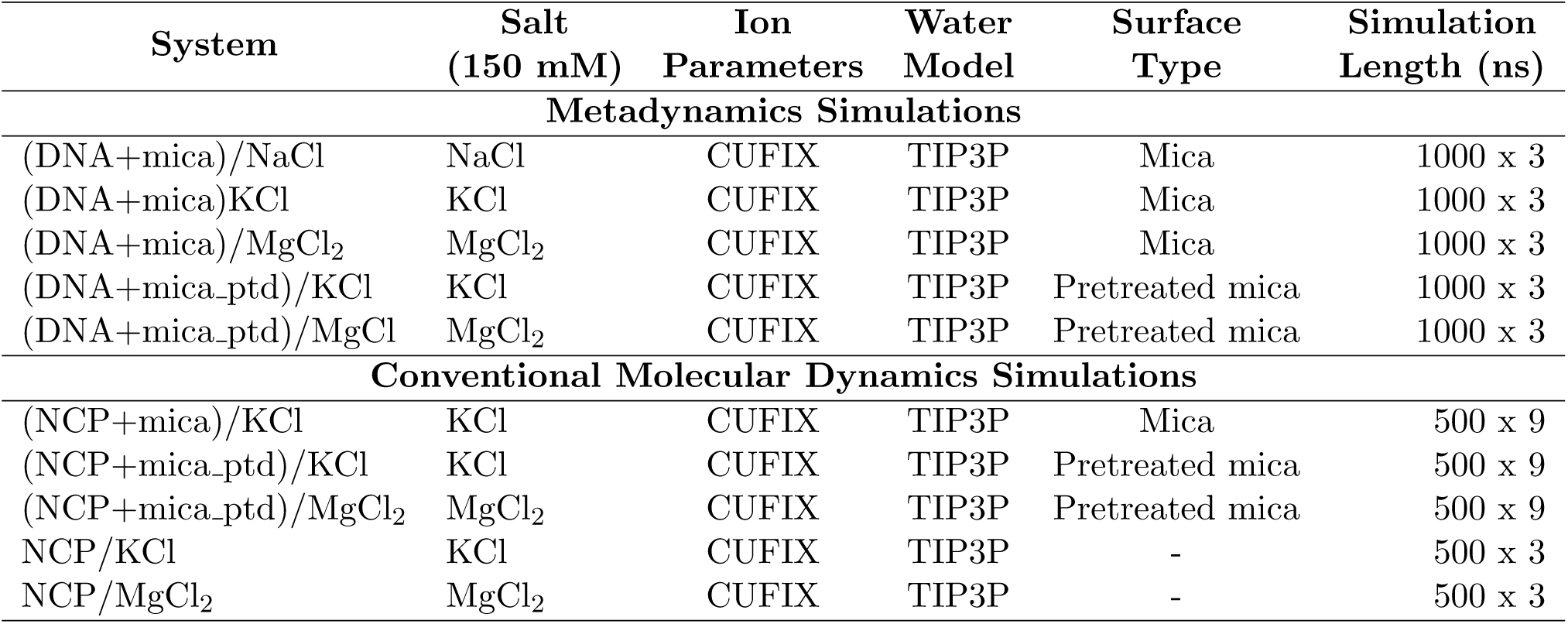
Summary of metadynamics and conventional molecular dynamics simulations performed.

| System | Salt<br>(150 mM) | Ion<br>Parameters | Water<br>Model | Surface<br>Type | Simulation<br>Length (ns) |
| --- | --- | --- | --- | --- | --- |
| **Metadynamics Simulations** |  |  |  |  |  |
| (DNA+mica)/NaCl | NaCl | CUFIX | TIP3P | Mica | 1000 x 3 |
| (DNA+mica)KCl | KCl | CUFIX | TIP3P | Mica | 1000 x 3 |
| (DNA+mica)/MgCl<sub>2</sub> | MgCl<sub>2</sub> | CUFIX | TIP3P | Mica | 1000 x 3 |
| (DNA+mica_ptd)/KCl | KCl | CUFIX | TIP3P | Pretreated mica | 1000 x 3 |
| (DNA+mica_ptd)/MgCl | MgCl<sub>2</sub> | CUFIX | TIP3P | Pretreated mica | 1000 x 3 |
| **Conventional Molecular Dynamics Simulations** |  |  |  |  |  |
| (NCP+mica)/KCl | KCl | CUFIX | TIP3P | Mica | 500 x 9 |
| (NCP+mica_ptd)/KCl | KCl | CUFIX | TIP3P | Pretreated mica | 500 x 9 |
| (NCP+mica_ptd)/MgCl<sub>2</sub> | MgCl<sub>2</sub> | CUFIX | TIP3P | Pretreated mica | 500 x 9 |
| NCP/KCl | KCl | CUFIX | TIP3P | - | 500 x 3 |
| NCP/MgCl<sub>2</sub> | MgCl<sub>2</sub> | CUFIX | TIP3P | - | 500 x 3 |

The TIP3P-FB (Force Balance)^31^ and TIP4P-FB water models^31^ with CUFIX ion parameters were also tested to observe any dependence of the water model on binding free energy. Both the hexahydrated Mg^2+^ ion model and the regular Mg^2+^ ion model without bonded water (labeled as MgCl_2_ std) were also tested for binding affinity. Additionally, three other systems were prepared at 25, 50, and 75 mM concentrations for both KCl and MgCl_2_ with the regular mica surface to study the effects of ion concentration on binding affinity. The TIP3P water model with CUFIX ion parameters was used for these systems. A summary of these metadynamics simulations is provided in Table S1.

All systems were energy minimized, heated, and equilibrated prior to metadynamics simulations as described in the molecular dynamics simulations section below. The vertical distance between the center of mass of the DNA and the center of mass of the surface oxygen atoms of the mica surface was used as the collective variable. Well-tempered metadynamics simulations were performed with a Gaussian hill of height 0.2 deposited every 2 ps and a metadynamics bias temperature of 1500 K. ^32–34^ These calculations were performed with the NAMD colvars module,^35^ and each simulation was run for 1 µs, with either three or one copies of each.

### Molecular Dynamics Simulations

The nucleosome crystal structure 5NL0^36^ was used as the initial structure. Modeller via Chimera^37^ was used to build the missing loops and tails for each histone. Only the 147 base pairs of nucleosomal DNA was retained, and the linker histone was removed.

A large double-layer surface of muscovite mica (140 Å *×* 153 Å *×* 20 Å) was generated using CHARMM-GUI^22^ for the nucleosome simulations. Then, a 180 Å-thick water box was constructed on top of the mica surface using VMD. Initially, the nucleosome was placed approximately 80 Å away from the mica surface. Three different systems were set up to illustrate the effects of ion type in solution and the ions adsorbed on the mica surface on nucleosome binding. The magnesium pre-treated mica surface (mica ptd) was obtained by randomly replacing one-fourth of the potassium ions adsorbed on the regular mica surface with magnesium ions. The number of replaced ions was calculated using the fraction of magnesium ions adsorbed on the mica surface during a 4 µs equilibrium simulation of mica with MgCl_2_ for a smaller mica surface. A representation of the pre-treated and regular mica surfaces is shown in Figure 1.

**Figure 1:**
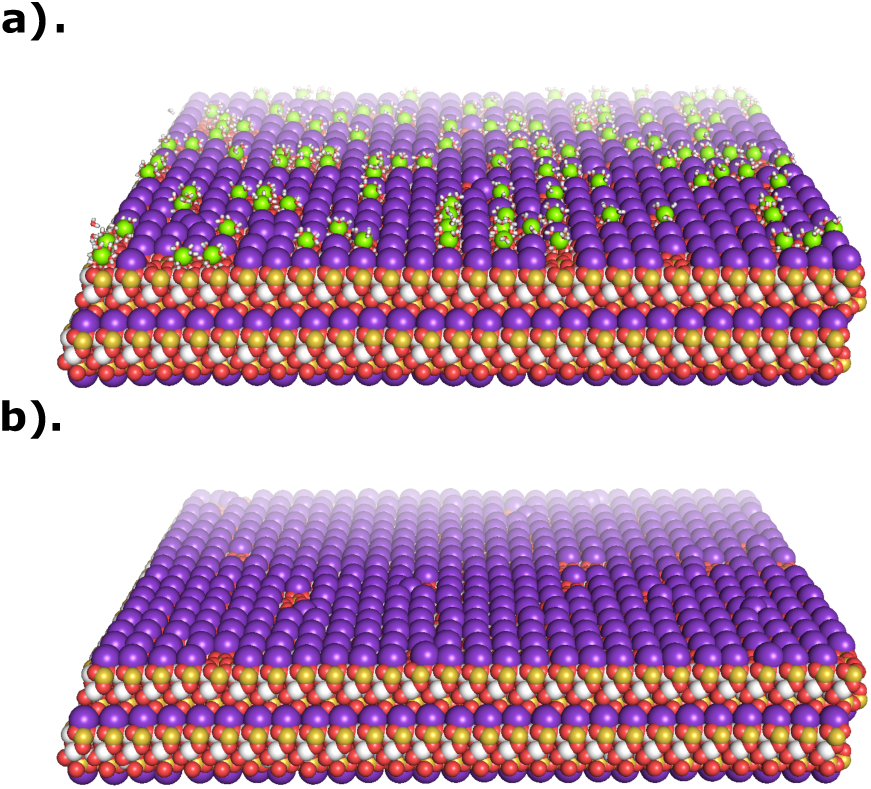
Double-layered mica surfaces after the first 50 ns of equilibrium simulations. (a) Magnesium ion pre-treated mica surface (mica ptd), where one-fourth of potassium ions (purple spheres) on the top layer were randomly replaced with magnesium ions (green spheres). Water molecules bound to hexa-hydrated magnesium ions are shown in stick representation. (b) Regular mica surface with adsorbed potassium ions. Nucleosome and solvent are not shown for clarity. Figure generated using PyMOL molecular visualization software.^45^

The first system was prepared by solvating the nucleosome in 150 mM KCl with a regular mica surface where potassium ions were adsorbed to the mica surface ((NCP+mica)/KCl). The second system was prepared by solvating the nucleosome in 150 mM KCl solution with a pre-treated mica surface ((NCP+mica ptd)/KCl). The third system was prepared by solvating the nucleosome in 150 mM MgCl_2_ solution, where the mica surface was pre-treated using the magnesium ions present in the solution with the same ratio as mentioned above ((NCP+mica ptd)/MgCl_2_).

As in the DNA/mica metadynamics simulations, the protein and DNA force field parameters were CHARMM36m^38^ and CHARMM36,^39^ respectively. The mica surface was parameterized via the INTERFACE force field,^26^ and the ions were treated with the CUFIX parameters developed for the CHARMM force field.^27^ Hexa-hydrated Mg^2+^ ion model ([Mg(H_2_O)_6_]^2+^) was used for the pre-treated mica surface and the Mg^2+^ ions in solution.

Each system was minimized twice for 5000 steps: first with a 10 kcal/mol/Å^2^ harmonic restraint applied to all heavy atoms of the solute and mica surface, and then without restraints. Systems were then gradually heated from 10 K to 300 K with restraints, which were subsequently reduced in an isothermal-isobaric ensemble. All simulations were performed in the NPT ensemble at 300 K and 1 bar pressure with a Langevin thermostat and a Langevin piston barostat^40,41^ within NAMD. The simulation box was periodic in all dimensions and was allowed to fluctuate only in the z direction to maintain constant pressure. The van der Waals forces were cut off at 12 Å, and electrostatic interactions were treated with the Particle Mesh Ewald method (PME).^42^ A time step of 4 fs was used with hydrogen mass repartitioning.^43^ All simulations were performed using NAMD 2.14.^35^ The bottom mica layer was restrained using a 3 kcal/mol harmonic restraint during equilibrium simulations to mimic bulk effect as per Köohler and coworkers.^44^

After relaxing the systems, 50 ns of equilibrium simulations were performed. Then, triplicates of steered molecular dynamics simulations were performed for 50 ns for each system using the NAMD colvars module. The vertical distance between the center of mass of the nucleosome core protein and the center of mass of oxygen atoms on the mica surface was used as the reaction coordinate for steered molecular dynamics simulations. These simulations were used to obtain three different starting structures, where the center of mass distance between the nucleosome and mica surface was 45, 50, and 55 Å. Triplicates of equilibrium simulations were performed at each window for 500 ns,totalling 4.5 *µ*s for each NCP+mica systems and were preceded by minimization and heating. A harmonic potential was applied to the histone core atoms along the vertical direction to keep the nucleosome at the initial position for the first 25 ns of the production run.

Two additional systems were set up by solvating the nucleosome in 150 mM KCl and MgCl_2_ solutions (NCP/KCl and NCP/MgCl_2_) to compare the nucleosome structure in the absence of a mica surface. Triplicates of equilibrium simulations were performed for 500 ns, followed by minimization, heating, and relaxation.

### Simulation Analyses

The root mean square deviation (RMSD) of the core protein and the DNA backbone was calculated after fitting to the initial conformation using the AMBER CPPTRAJ tool.^46^

Residue root mean square fluctuations (RMSFs), end-to-end DNA distance, and radius of gyration of histone tails were also calculated using the CPPTRAJ tool.

Angle and contact analysis was performed using MDAnalysis^47,48^ and custom Python scripts. Two angles were defined relative to the mica surface: the side angle and the forward angle. The side angle was calculated using the angle between the vector passing through the center of mass (COM) of the histone core residues and the COM of half of the DNA to the plane perpendicular to the mica surface. The forward angle was calculated using the vector passing through the COM of the histone core residues and the dyad.

Cluster analysis was performed with k-means clustering, and the number of clusters was determined based on the Silhouette coefficient value. The vertical distance from any heavy nucleosome atom to the center of mass of oxygen atoms on the mica surface was used to calculate the number of contacts with the mica surface. The residues of the N- and C-terminal tails were included in the calculation of the mica-tail contacts, with 5 Å used as the contact distance. For DNA-mica contacts, 8.2 Å was used as the contact distance for the two systems with Mg^2+^ pre-treated mica surfaces, while 6.5 Å was used as the contact distance for the KCl system with a regular mica surface. Different distances were used because the hexahydrated magnesium ions adsorb slightly above the mica surface, positioning the nucleosomes slightly away from the oxygen layer of the mica surface compared to the regular mica surface with potassium ions. The center-of-mass distance between the adsorbed cations on the mica surface and the oxygen atoms on the mica surface was added to the distance of 5 Å when calculating DNA-mica contacts, as DNA interacts with the adsorbed cations on the mica surface. The first 100 ns of simulation data were excluded from the analysis except for the time history analysis.

## Results

### Solvent Effects on DNA/Mica Interactions

Multiple all-atom force fields have been developed for water, monovalent, and divalent ions, each of which can uniquely influence solute/solute and solute/solvent interactions.^49,50^ To identify the solvent conditions and force field parameters most likely to create stable nucleosome/mica interfaces, we first quantified the free energy of DNA/mica interactions through a series of well-tempered metadynamics simulations.^51^ These simulations used the vertical distance between the center of mass of a Drew-Dickerson DNA molecule and the center of mass of oxygen atoms on the mica surface as a reaction coordinate (Figure S1).

Simulations were carried out in a 150 mM NaCl, 150 mM KCl, and 150 mM MgCl_2_ environment (Table 1), where the hexahydrated magnesium model was used. Simulations were also performed for two variations of the mica surface, one in which potassium ions coated the surface, as generated with CHARMM-GUI, and a “pre-treated” surface (which we denote as mica ptd) in which potassium ions were allowed to exchange with bulk magnesium during a 4 µs equilibrium simulation. This pre-treated surface resulted in a ratio of approximately one magnesium per four potassium ions on the surface.

Metadynamics results showed that solution conditions strongly influence the thermodynamics of DNA binding to the mica surfaces (Figures 2). In TIP3P water, DNA bound weakly to mica in NaCl, with a minimum binding free energy of -5.6 *±* 0.8 kcal/mol. Replacement of NaCl with KCl created a stronger DNA/binding interface, with a binding free energy of -7.3 *±* 0.7 kcal/mol, while when MgCl_2_ was present in solution, the DNA binding affinity was weak and similar to that of NaCl (-5.3 *±* 0.4 kcal/mol). The latter was particularly surprising in light of experiments that have indicated strong DNA binding in the presence of divalent ions.^52–54^ This result inspired the use of magnesium-pretreated surfaces, which indeed created a much stronger binding affinity of -9.2 *±* 0.5 kcal/mol with bulk magnesium. The binding affinity further increased for the KCl system with the pretreated mica surface (-11.4 *±* 0.7 kcal/mol), which gave the minimum binding free energy of the five systems.

**Figure 2:**
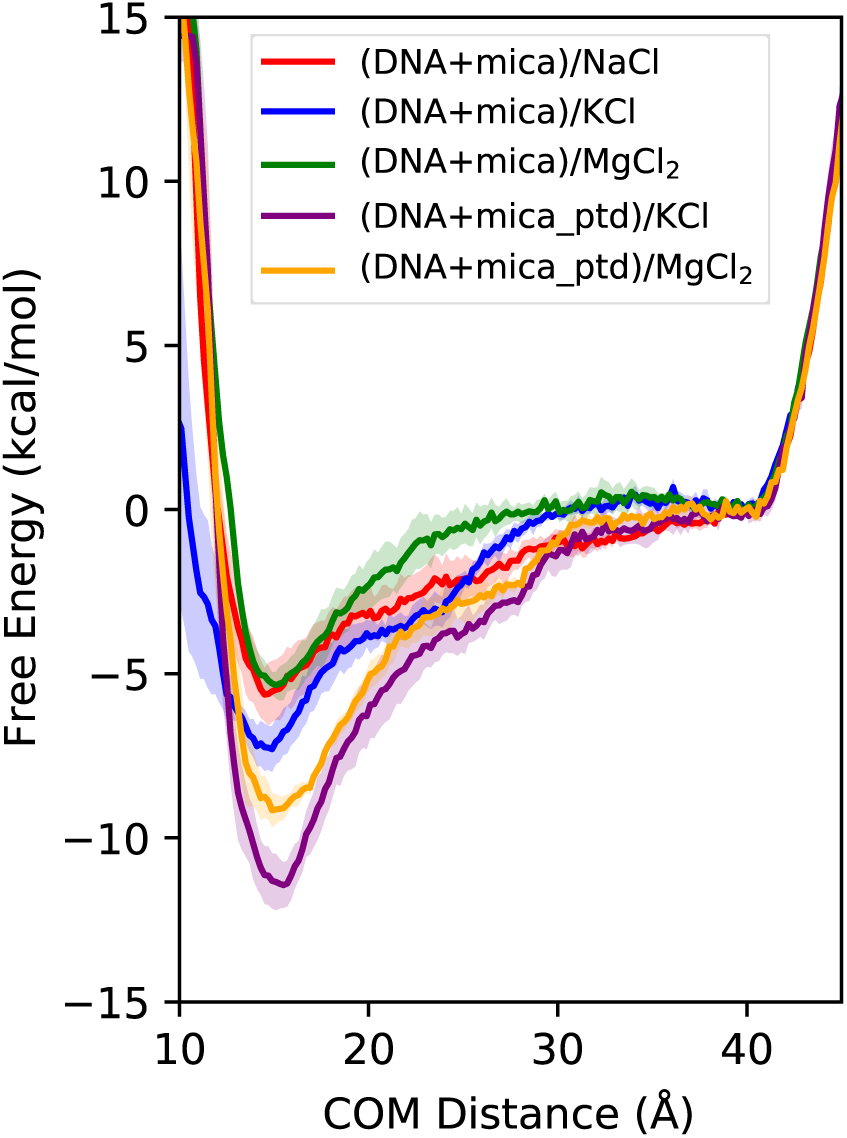
Free energy profiles of DNA binding to the mica surface in the presence of NaCl, KCl, and MgCl_2_, on regular mica surface and in the presence of MgCl_2_ on pre-treated mica surface with TIP3P water model.

To determine whether other solvent force field effects, such as the water or the magnesium ion models, affected DNA binding to mica surfaces, additional metadynamics calculations were performed. Simulations with the TIP3P-FB and TIP4P-FB water models showed little difference compared to those of TIP3P (Figure S2), and therefore we proceeded with the TIP3P model for the remainder of our simulations. To check whether the hexahydrated magnesium ion model prevents DNA binding to the mica surface due to its solvation shell, the binding free energy was calculated with the regular Mg^2+^ ion model (Figure S3). However, the system with a hexahydrated Mg^2+^ model provided a stronger DNA binding affinity to the mica surface, and therefore we kept the hexahydrated Mg^2+^ ion model for the rest of our simulations. Furthermore, we compared the effect of salt concentration on DNA binding affinity to mica surfaces and observed that the binding affinity can be fine-tuned by salt concentration (Figure S4). The highest binding affinity was observed at 75 mM salt concentration with both KCl and MgCl_2_ and both systems showed similar affinities at this concentration. However, to determine the nucleosome binding affinity on the mica surface under physiological conditions, we kept the 150 mM salt concentration for the remainder of our nucleosome/mica simulations.

Visual inspection of the metadynamics trajectories suggested that the orientation of DNA relative to the mica was affected by the solvent environment. To test this, we calculated the angle distribution of the DNA major axis relative to the mica surface as a function of the center of mass distance to the mica surface (Figure S5). Systems with DNA in the KCl solution showed narrow angle distributions as the DNA approached the surface, with the DNA mostly staying parallel to the mica surface. In NaCl and MgCl_2_ with a regular mica surface, there was a wider angle distribution far from the mica surface with more DNA orientation sampling. However, replacement of the regular mica surface with a Mg^2+^ pretreated mica surface in the MgCl_2_ system limited this distribution as the DNA sampled fewer states far from the surface.

### Ionic Conditions Influence Nucleosomes Binding to Mica Surfaces

Based on the observation that DNA preferably binds to mica surfaces with KCl in solution or Mg^2+^ pretreated mica surfaces, we set up three different nucleosome plus mica systems: I. a nucleosome in KCl with a regular mica surface ((NCP+mica)/KCl), II. a nucleosome in KCl with a pretreated mica surface ((NCP + mica ptd)/KCl), and III. a nucleosome in MgCl_2_ with a pretreated mica surface ((NCP +mica ptd)/MgCl_2_. The TIP3P water model with CUFIX ion parameters was used for each simulation. Three starting structures were selected for each system on the basis of the vertical distance between the center-of-mass of the nucleosome core and the center-of-mass of the oxygen atoms on the mica surface. Equilibrium molecular dynamics simulations were performed in systems where the nucleosome was initially located 45, 50, and 55 Å away from the mica surface, and for each starting distance, three 500 ns simulations were performed, for a total of 4.5 µs of sampling per system. Different distances were chosen to sample the nucleosome binding process to mica surfaces, as it allowed us to observe whether nucleosomes move away from the surface when placed closer to it or move closer to it when placed far from the surface.

The center of mass distance between the nucleosome core and the mica surface was measured as a function of simulation time (Figure 3). In general, nucleosomes in KCl remained close to or moved closer to the regular and pre-treated mica surfaces, with the one exception being run 1 of the (nuc+mica)/KCl system initiated at a distance of 50 Å. Similarly, nucleosomes in MgCl_2_ with the pretreated mica surface stayed within a couple of angstroms of their initial distance or moved closer to the surface except in two simulations: runs 1 and 2 of the 55 Å window. In those two simulations, the nucleosomes moved further and did not make contact with the mica surface. In some simulations, the center-of-mass distance remained relatively constant despite changes in the nucleosome/surface interactions, mainly because nucleosomes could have the same vertical center-of-mass separation but different angle orientations, as discussed below.

**Figure 3:**
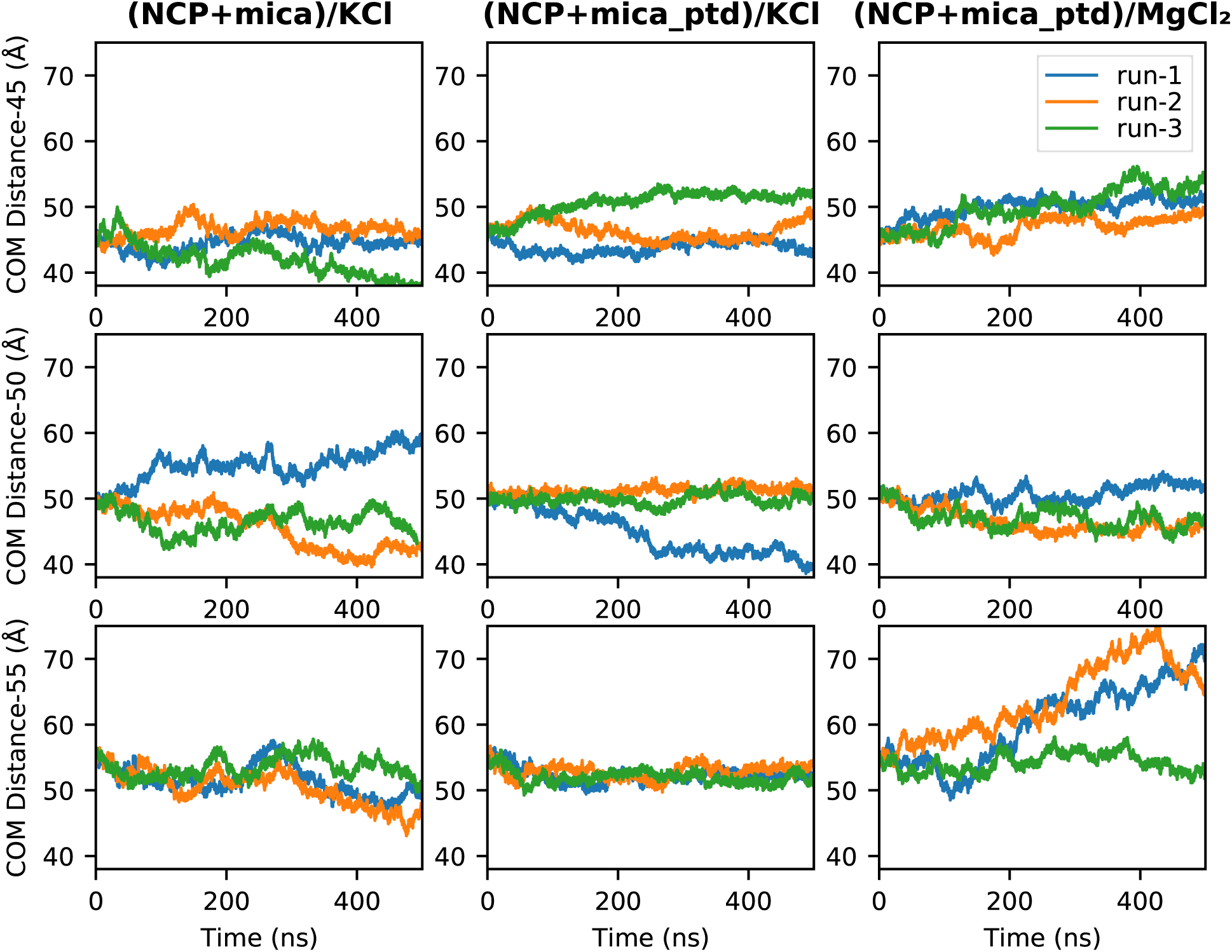
Center of mass (COM) distance of nucleosome core to the mica surface for three systems started at 45 Å, 50 Å, and 55 Å away from mica surface. Three replicates for each windows are shown in blue, orange and green color lines.

To quantify the orientation of the nucleosome relative to the mica surface, we calculated two angles between the nucleosomal plane and the mica surface, which we denote as the forward and side angles (Figure 4). Forward and side angles of zero degrees correspond to a nucleosome orientation that is parallel to the mica surface. Angle distributions indicate that the nucleosome can have different orientations in solution and also when contacting the mica surface. The nucleosomes in the KCl solution had a much more localized angle distribution compared to the nucleosome in MgCl_2_, which sampled a wide range of COM distances with different orientations when not in contact with the mica surface.

**Figure 4:**
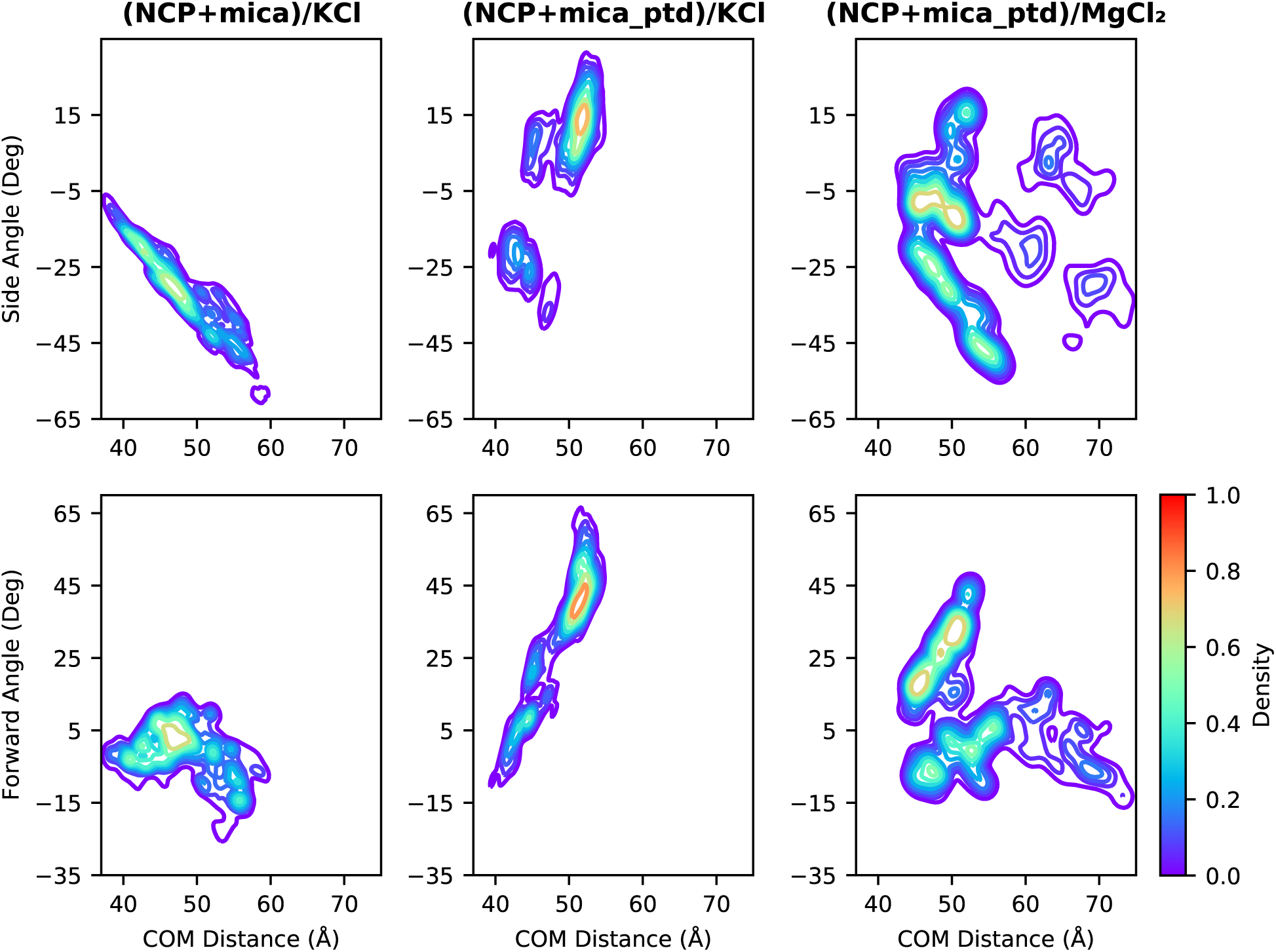
Forward and side angle as a function of COM distance between the histone core residues and mica surface.

A K-means cluster analysis used the COM distances and angles to determine the dominant nucleosome states near the mica surfaces. Three main clusters were observed for the (NCP+mica)/KCl and (NCP+mica ptd)/KCl systems, while eight clusters were observed for (NCP+mica ptd)/MgCl_2_ system, which represent the major distinct orientations of nucleosomes relative to the mica surface (Figure 5). In most dominant orientations, only a fraction of DNA interacted with the mica surface, as there were significant forward or side tilt angles. The exception to this was the second orientation of the (NCP+mica)/KCl system, which is mainly parallel to the mica surface and accounts for 30% of the simulation time. An increased forward angle was observed for the (NCP+mica ptd)/KCl system as the nucleosome interacted with the mica surface through the off-dyad side of DNA (orientation 1 and 2 of NCP+mica ptd)/KCl). The (NCP+mica ptd)/MgCl_2_ system had both parallel and tilted interaction modes and provided eight clusters and a wide range of COM distances when unbound to mica.

**Figure 5:**
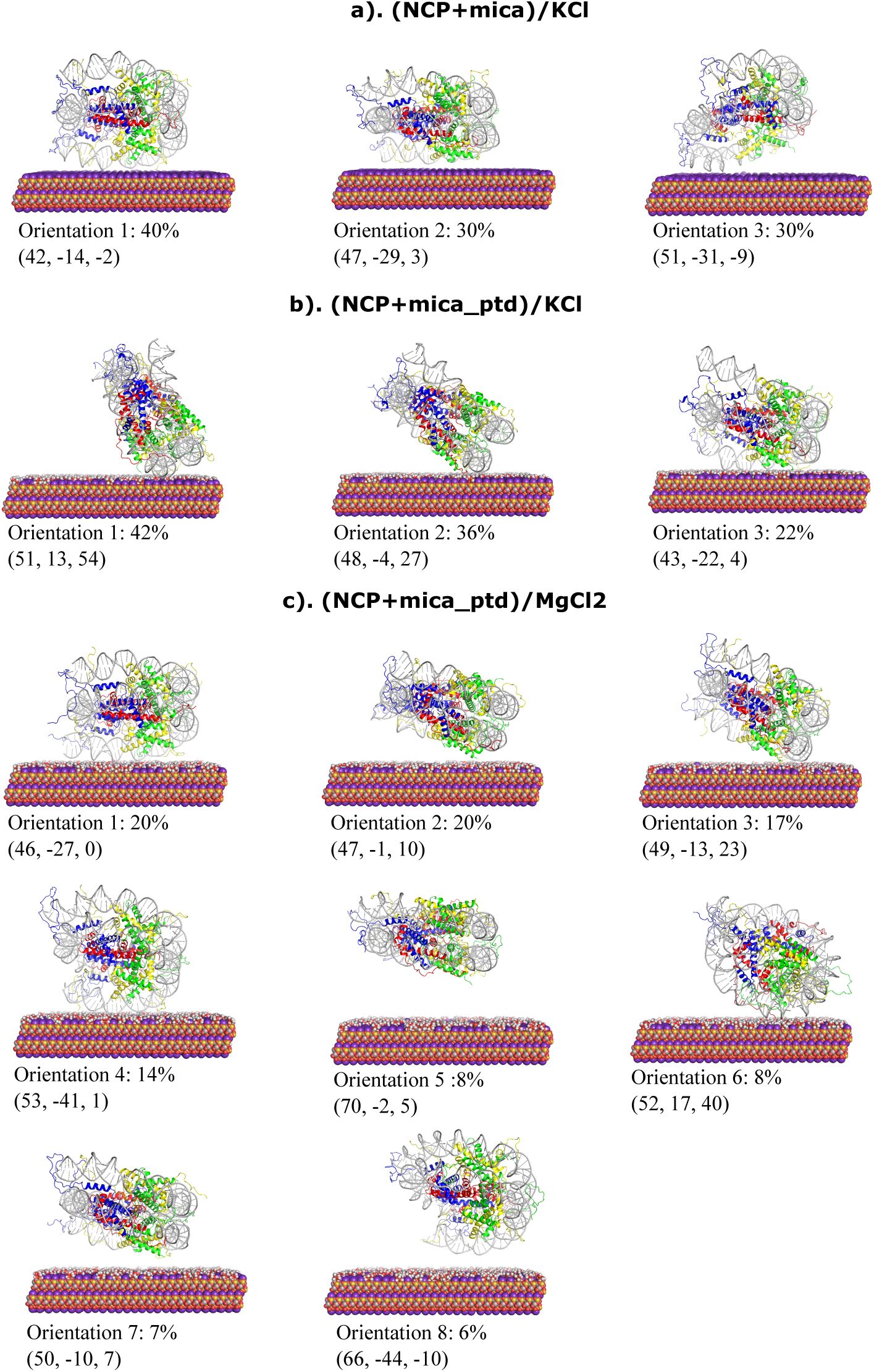
Representative nucleosome orientation snapshots obtained based on 3D-cluster analysis. Percentage of each cluster is labeled. The COM distance (Å), Side, and Forward angles (Deg) corresponding to each conformation are shown within the parenthesis. Histone chains, H3, H4, H2A, and H2B are colored in blue, red, yellow, and green colors respectively. DNA is shown in grey color. Images were generated with PyMOL.^45^

To determine which nucleosome components were primarily responsible for stabilizing nucleosome/mica interactions, the number of contacts between the mica surface and nucleosomal DNA, histone tails, and histone core were calculated. The average number of contacts for each constituent shows that in each case and simulation starting distance, the nucleosomal DNA made the majority of contacts on the mica surface, followed by the histone tails, with the histone core making few contacts, if any (Figure 6). This effect was the most pronounced in the KCl with pre-treated mica surface simulations in which the DNA made approximately 37 contacts to the mica surface regardless of the starting configuration, followed by only 5 contacts of the histone tails. In both KCl with regular mica and MgCl_2_ with pre-treated mica simulations, the number of DNA contacts was reduced by approximately half. In each case, the negatively charged nucleosomal DNA interacted with the mica via positively charged cations adsorbed on the surface, while the histone tails interacted via cationic residues (lysine and arginine), which made direct contacts with the negatively charged mica surface.

**Figure 6:**
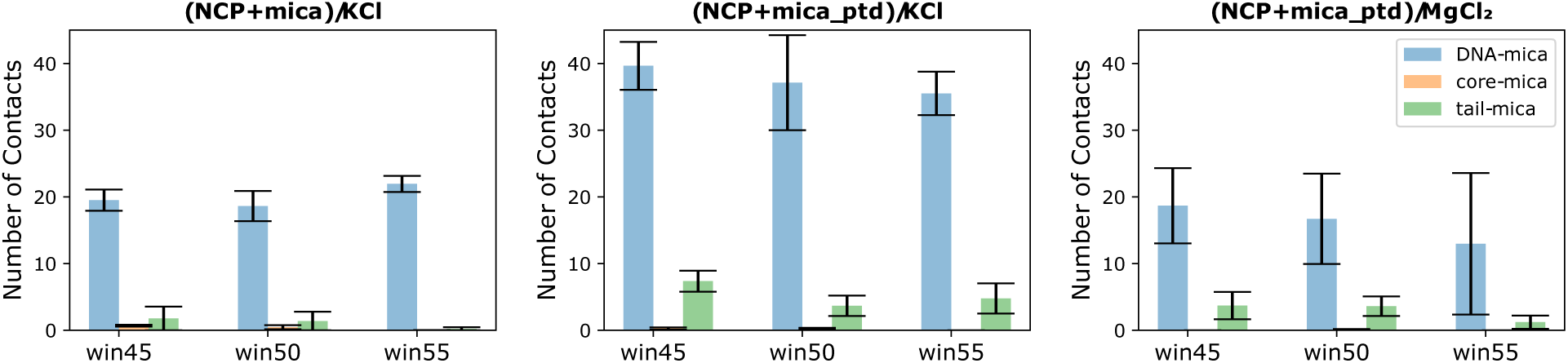
Average number of DNA-mica, histone tail-mica and histone core-mica contacts for each system at different windows.

In addition, solution cations can directly affect nucleosome/mica binding. To examine this, we calculated the charge of the cations within 5 Å of the nucleosomal DNA and plotted it as a function of the center of mass distance between the nucleosome core and mica surface (Figure 7). In general, the cation charge around the nucleosomes in MgCl_2_ was significantly higher than that around those in KCl solution. The total nucleosome charge in our simulations was -144*e*, and therefore with an average cation charge of approximately +115*e* surrounding the nucleosome in the (NCP+mica ptd)/MgCl_2_ system there was a significant charge neutralization by Mg^2+^ ions in solution. This neutralization may hinder the adsorption of nucleosomal DNA onto the Mg^2+^ pre-treated mica surface due to repulsive forces between the cation layers. In contrast, for both KCl systems, the cation charge around the DNA was only around +80*e*, indicating that the nucleosome maintained more of its effective negative charge, thus allowing increased attraction for adsorbed cations on the mica surface.

**Figure 7:**
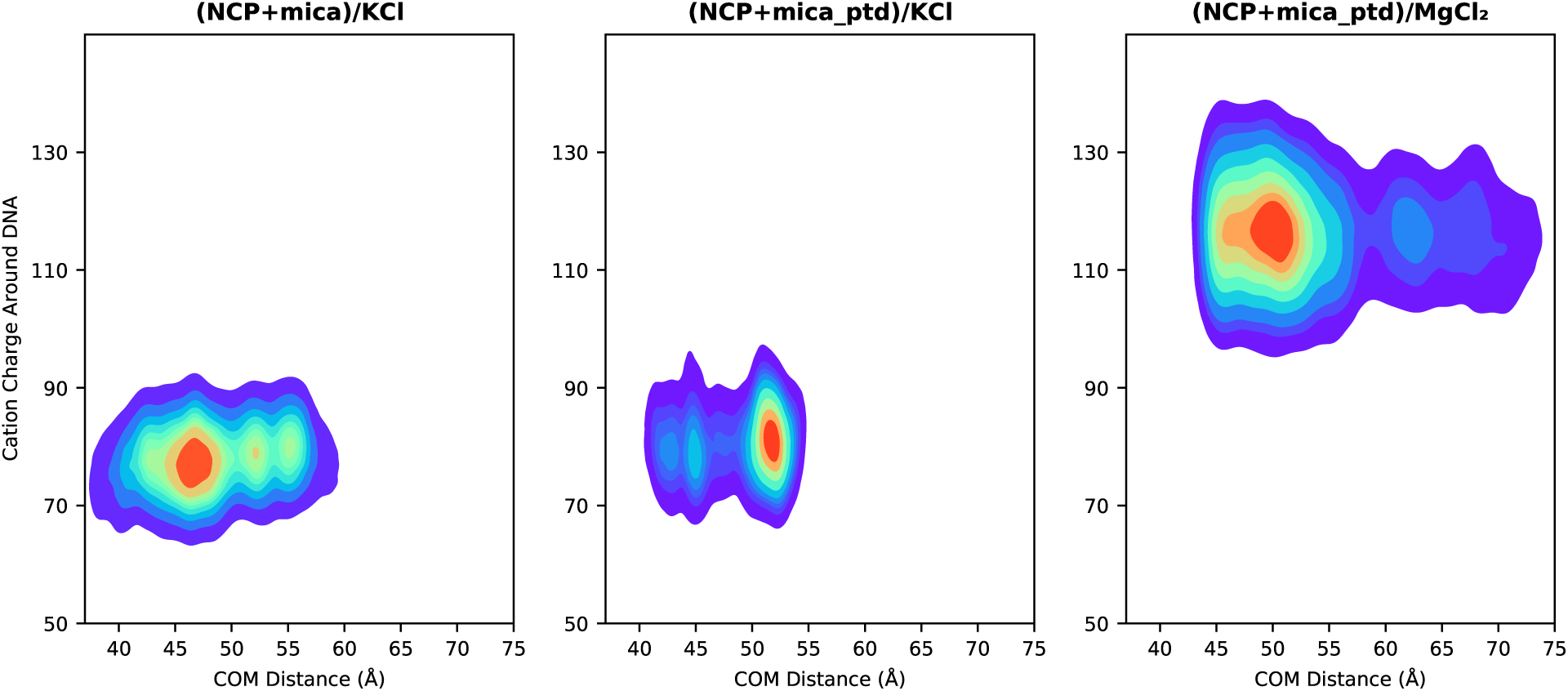
Cation charge within 5 Å of nucleosomal DNA as a function of minimum z-distance from any heavy atom of DNA to mica surface.

### Nucleosomal Structural Changes upon Mica Binding

Having established that nucleosome/mica binding is mediated by the cations adsorbed to the mica surface and depends on the ion type in solution, we sought to determine how this process affects nucleosomal structures and dynamics. The root mean square deviations (RMSDs) of the histone core residues did not show large-scale structural changes upon binding to the mica surface, with RMSD values ranging from 2-4 Å and the highest and lowest values in the (NCP+mica ptd)/KCl and (NCP+mica ptd)/MgCl_2_ systems respectively (Figure S6). Non-zero RMSD values were observed at the beginning of these simulations, as these were performed after 50 ns of equilibrium simulation and another 50 ns of steered MD simulations for systems with the mica surface, and the RMSDs were calculated using the initial nucleosome structure. Although relatively small, we note that these RMSD values are higher than the RMSD values for nucleosomes in solution, which ranged from 1.0-1.5 Å (Figure S7 top row), indicating that mica binding causes some degree of subtle structural rearrangement of the core structure on the hundreds of nanoseconds timescale beyond standard thermal fluctuations. Similar trends were observed for DNA RMSDs, which were also typically higher in the mica surface system than without it by 1-3 Å (Figures S8 and bottom row of S7).

One of the most studied nucleosome dynamic modes is DNA breathing, in which one or both ends of DNA separate from the histone core.^55,56^ Although no significant DNA breathing was observed on the timescales simulated here, the ion type and the distance to the mica surface affected the DNA end-to-end distance, which is indicative of the DNA’s propensity to unwrap (Figure 8, Figure S9). In solution, DNA sampled end-to-end distances of 58 to 85 Å in the KCl environment, which were reduced to 55 to 80 Å in MgCl_2_, with both systems exhibiting most likely distances of around 70 Å (Figure S12). These reduced distances in the Mg^2+^ ion solution suggest an increased stability of the ions in DNA compared to the K^+^ ions. This distribution was narrowed with slightly more DNA compaction in the (NCP+mica)/KCl case, where the most likely DNA end separation was 65-68 Å regardless of the initial configuration. However, in the (NCP +mica ptd)/KCl simulations, DNA was observed to be in more open states, especially in the windows that began closer to the mica surface, with the most likely end-to-end DNA distances being 72 and 76 Å for the 50 Å and 45 Å windows, respectively. This is attributed to the increase in the number of contacts made with the pretreated mica surface in this system, as discussed above. For the (NCP+mica ptd)/MgCl_2_ simulations there were also increased end-to-end distances relative to both solution states, although the distributions were not as extended as the (NCP+mica ptd)/KCl systems.

**Figure 8:**
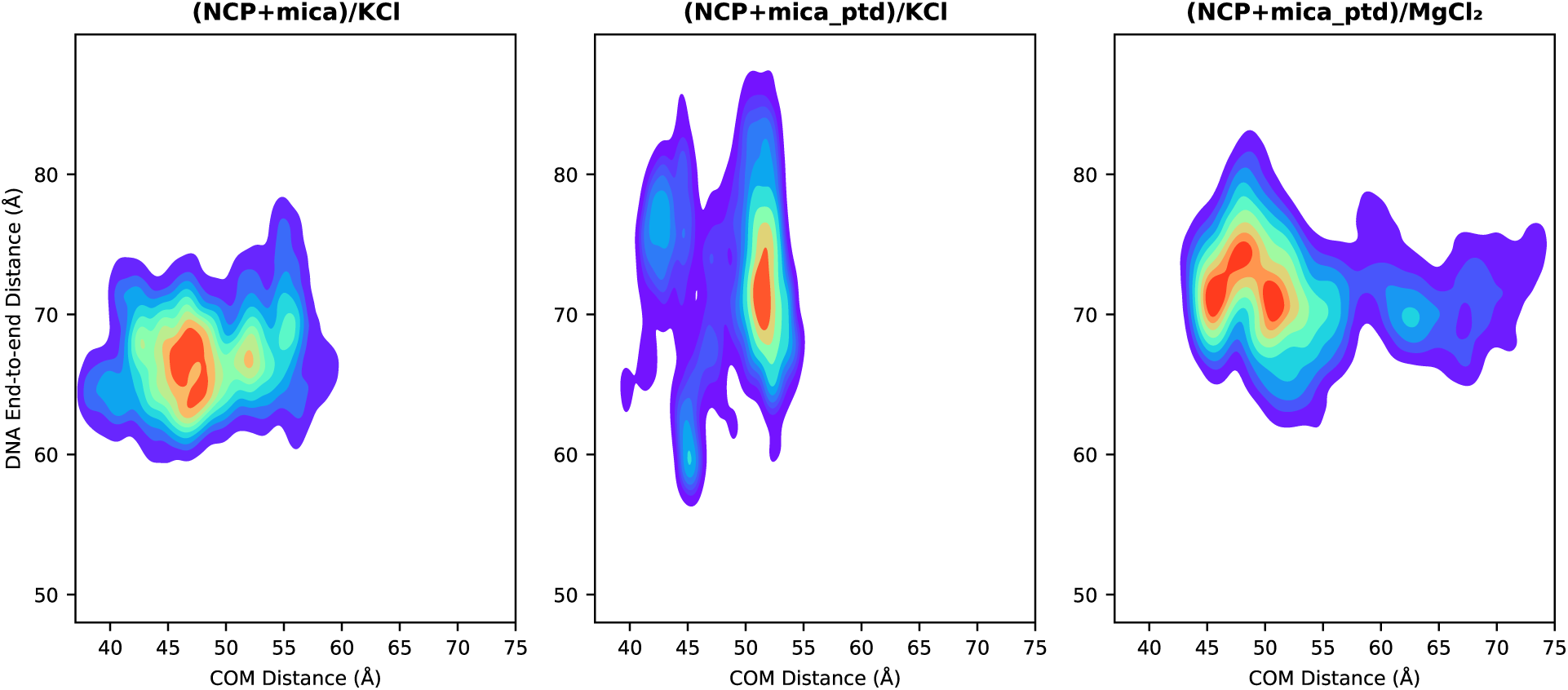
DNA end to end distance vs the COM Z-distance from histone core to mica surface.

The root mean square fluctuations (RMSF) of histone proteins were calculated to identify the regions with the highest fluctuations (Figure S10). No significant differences in the fluctuations in histone core protein residues were observed between the three nucleosome and mica systems. However, an overall reduced fluctuation was observed in the histone tail regions compared to those observed in solution, indicating that the dynamics of histone tails is altered in the presence of mica. To characterize the structures of these disordered regions, we computed the radius of gyration and the minimum tail/mica surface distance for each histone tail. The radius of gyration was plotted against the minimum distance from any heavy atom of the histone tail to the mica surface, allowing us to differentiate between the behavior of the tail when the tails are closer to the mica surface versus when they are further away (Figure 9).

Of particular note, in each system, the H2A tails made contact with the mica surface and, in doing so, adopted more extended states with increased radii of gyration. For example, in each system the most likely radius of gyration for an H2A tail in contact with the mica surface was approximately 15 Å, whereas when far from the surface with tail/mica distances greater than 40 Å the most likely radius of gyration was typically of the order of 5-10 Å. This was consistent with the visualization of the trajectories, which showed interactions between the N-terminal H2A tails and the mica surface in each of the three systems. Similarly, the H4 tails made contact with the mica surfaces in the pre-treated KCl and MgCl_2_ systems, which was associated with relatively elongated states of 16-19 Å. Interestingly, the H3 tails, which are the longest of the histone tails and have been shown to adopt an ensemble of states in solution,^57,58^ did not come into direct contact with the mica surfaces in any of the simulations.

**Figure 9:**
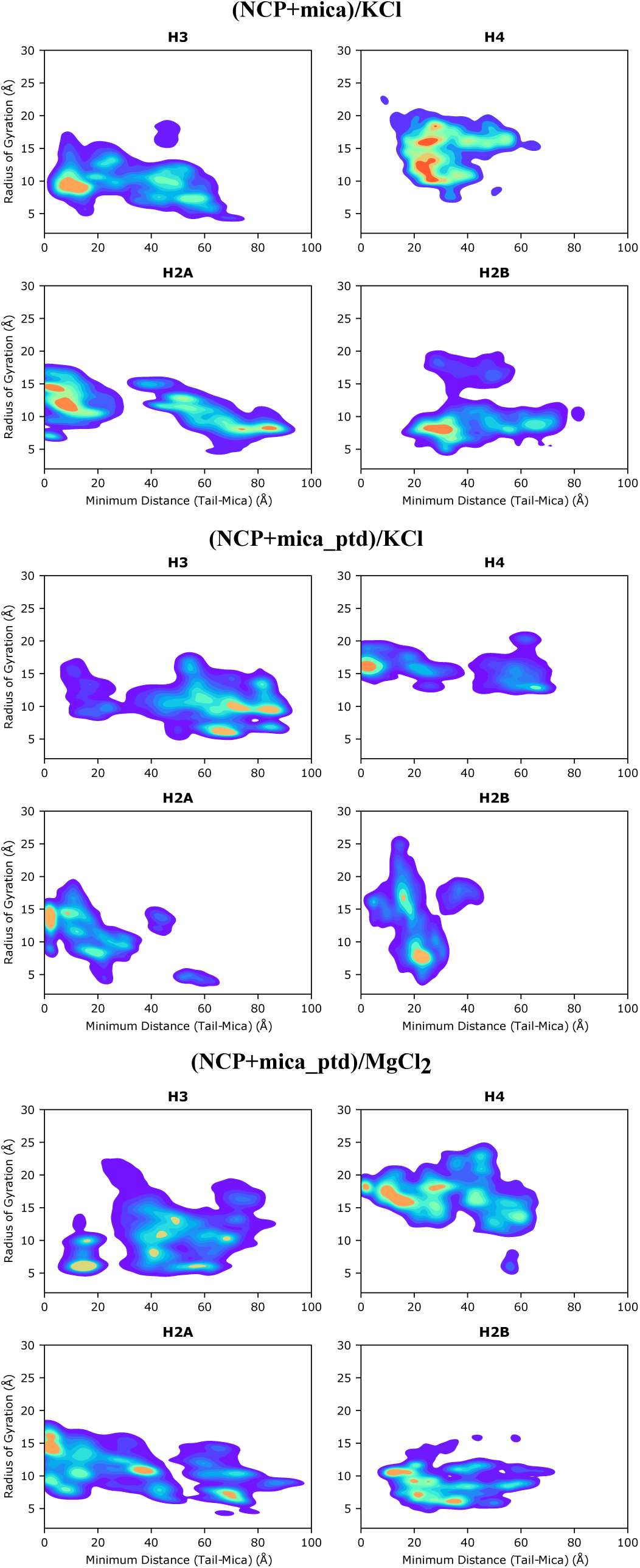
Radius of gyration of histone tails vs the minimum distance from any heavy atom of histone tail to the mica surface.

## Discussion

Here, we performed a series of atomistic molecular dynamics simulations to investigate nucleosome binding to a mica surface under different ionic environments. We first identified optimal force field parameters and ionic conditions through metadynamics simulations of DNA with both regular and pre-treated mica surfaces in various ionic environments. Our results showed that DNA-mica binding affinity can be finely tuned by the choice of counter ions and surface modifications. The lowest binding free energy was observed with mica surfaces pretreated with Mg^2+^ ions in a KCl solution, followed by MgCl_2_ in solution with a pre-treated mica surface. These results were in agreement with previous experiments which had shown that divalent cations bind DNA to mica surfaces more effectively than monovalent ions due to their higher charge density.^54,59^ An AFM study showed that the DNA binding free energy to mica was lowest with MgCl_2_ in solution compared to NaCl or KCl at 1 mM concentration, however the binding free energy increased as the MgCl_2_ concentration increased from 10 to 100 mM.^59^ Most AFM experiments typically use relatively low salt concentrations (*∼*10 mM).^20^

Our simulations of full-length nucleosomes indicated that nucleosomes bound more effectively to mica surfaces in KCl solutions, whether the mica was regular or pre-treated, at 150 mM ionic conditions. Nucleosome interactions with mica surfaces were consistent in all replicates for (NCP+mica)/KCl and (NCP+mica ptd)/KCl systems but occasionally moved away in the (NCP+mica ptd)/MgCl_2_ system. When nucleosomes interacted with the mica surface, their orientation varied. Nucleosomes adsorbed relatively flat in KCl solution with regular mica, while more tilted orientations were observed with pre-treated mica. These results indicated that nucleosome orientations with mica depended on the ionic environment and surface characteristics.

The highest number of contacts between nucleosomes and the mica surface was in the (NCP+mica ptd)/KCl system, while the lowest was in the (NCP+mica ptd)/MgCl_2_ system. In general, negatively charged DNA interacted with mica via cations adsorbed on the surface, and replacing some potassium ions with magnesium increased DNA-mica contacts compared to the (NCP+mica)/KCl system. However, nucleosomes did not bind effectively in the (NCP+mica ptd)/MgCl_2_ system. Our findings indicated that divalent magnesium ions exhibited a stronger affinity for nucleosomal DNA compared to monovalent potassium ions, which was consistent with previous simulations that showed a higher binding affinity of magnesium ions to DNA compared to monovalent ions.^60^ At 150 mM, magnesium ions effectively screened nucleosome charges, weakening interactions with mica-adsorbed cations. In contrast, KCl solutions provided less screening, allowing nucleosomal DNA to maintain its negative charge and interact effectively with the cations on mica’s surface.

Our simulations also showed structural changes in nucleosomal DNA upon mica interaction, including higher RMSD and RMSF values, and increased end-to-end distances in the (NCP+mica ptd)/KCl system compared to nucleosomes in solution. These measurements were compared to nucleosomes in solution without mica to isolate the effects of the mica surface. Additionally, it was observed that mica affected histone tail dynamics, with reduced RMSF values in the (NCP+mica ptd)/KCl system compared to nucleosomes in solution. The mica surface also impacted histone tail positioning and orientation, even in tails that did not directly interact with it. For example, H2A and H4 tails interacted with the surface and had increased radii of gyration near the surface due to their direct mica interactions.

Our study of three nucleosome systems with mica surfaces indicated that nucleosome structure depended on the ionic environment, and binding affinity could be fine-tuned with cations present both in solution and on the mica surface. AFM experiments require preserving the native properties of protein-nucleic acid complexes. Our findings demonstrated that regular mica with KCl solution at physiological concentrations facilitated effective nucleosome adsorption with minimal structural perturbation. However, 150 mM magnesium ions in solution heavily screened DNA, weakening interactions with mica-adsorbed cations. Overall, this study provides insights into nucleosome binding affinities and structural changes under different cationic conditions, highlighting the favorable binding of nucleosomes with monovalent potassium ions under physiological conditions.

## Acknowledgement

The authors thank Dr. Bibiana Onoa and Dr. Cèesar Díıaz-Celis for highly useful conversations which inspired this work. The authors also thank Wereszczynski group members, Dr. Lokesh Baweja for the useful comments and suggestions on this project and Dr. Dustin Woods for providing the initial nucleosome structure after modeling missing residues. This project was supported by the National Institutes of Health grant R35GM119647.

## Notes

### Competing Interest Statement

The authors have declared no competing interest.

