## Supporting Information for "The Influence of Ionic Environment on Nucleosome-Mica Interactions Revealed via Molecular Dynamics Simulations"

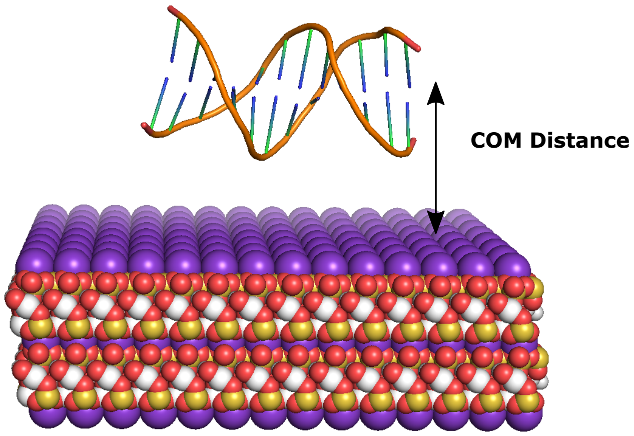


Figure S1: Representation of the reaction coordinate for the DNA+mica simulations, which is the the vertical distance between the center of mass of the DNA molecule and the center of mass of oxygen atoms on the mica surface.

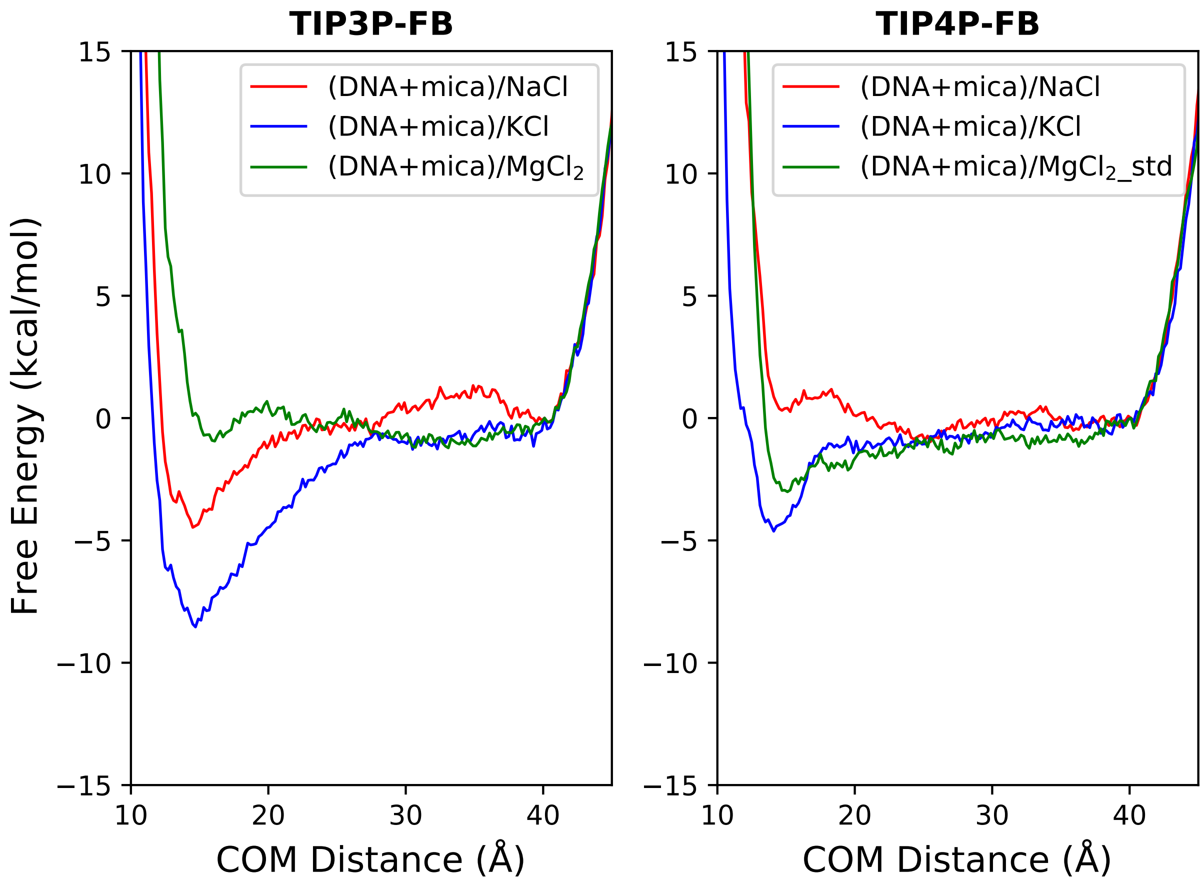


Figure S2: Free energy profiles of DNA binding to the mica surface in the presence of 150 mM NaCl, KCl, and $MgCl_2$, on regular mica surface with TIP3P-FB and TIP4P-FB water models. CUFIX ion parameters were used while regular $Mg^{2+}$ ion model was used with TIP4P-FB.

Table S1: Summary of metadynamics simulations to test the ion parameters, water models and salt concentrations.

| System | Solvent | Ion Parameters | Water Model | Surface Type | Simulation Length (ns) |
|---|---|---|---|---|---|
| (DNA+mica)/NaCl [TIP3P-FB] | NaCl | CUFIX | TIP3P-FB | Mica | 1000 |
| (DNA+mica)/KCl [TIP3P-FB] | KCl | CUFIX | TIP3P-FB | Mica | 1000 |
| (DNA+mica)/$MgCl_2$ [TIP3P-FB] | $MgCl_2$ | CUFIX | TIP3P-FB | Mica | 1000 |
| (DNA+mica)/NaCl [TIP4P-FB] | NaCl | CUFIX | TIP4P-FB | Mica | 1000 |
| (DNA+mica)/KCl [TIP4P-FB] | KCl | CUFIX | TIP4P-FB | Mica | 1000 |
| (DNA+mica)/$MgCl_2$_std [TIP4P-FB] | $MgCl_2$_std | CUFIX | TIP4P-FB | Mica | 1000 |
| (DNA+mica)/KCl [M] | KCl | CUFIX | TIP3P | Mica | 1000 |
| (DNA+mica)/$MgCl_2$ [M] | $MgCl_2$ | CUFIX | TIP3P | Mica | 1000 |
| (DNA+mica)/$MgCl_2$_std | $MgCl_2$_std | CUFIX | TIP3P | Mica | 1000 |

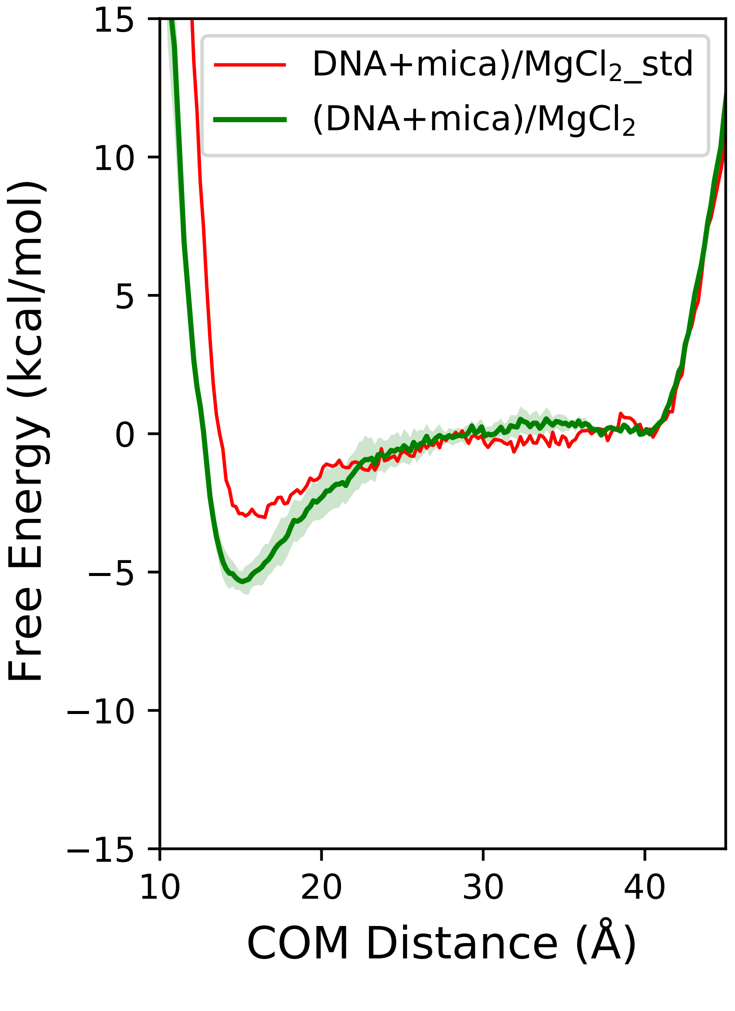


Figure S3: Free energy profiles of DNA binding to the mica surface with regular and hexa-hydrated $MgCl_2$ (150 mM), on regular mica surface with TIP3P water model.

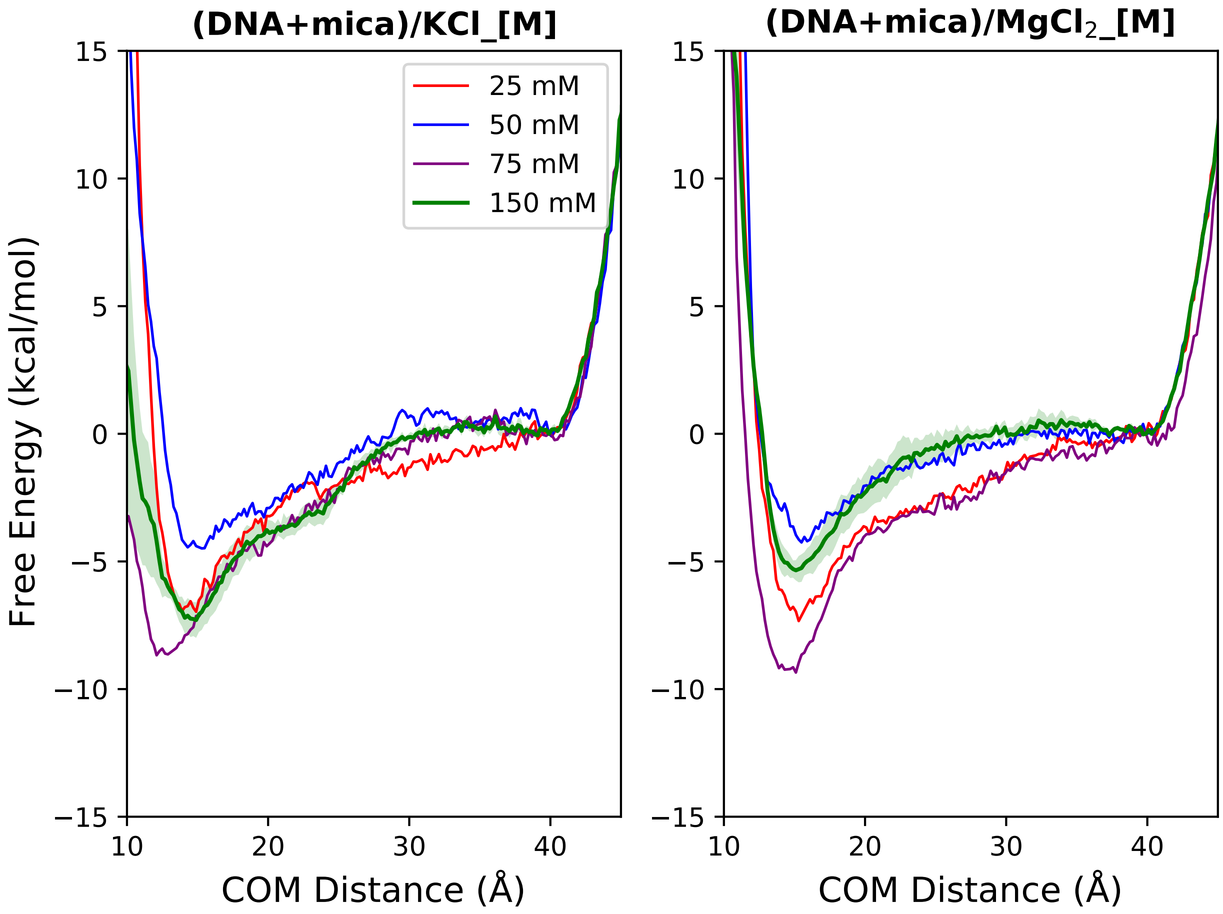


Figure S4: Free energy profiles of DNA binding to the mica surface in the presence of 25, 50, 75, 150 mM concentration of KCl, and $MgCl_2$, on regular mica surface. TIP3P water model with CUFIX ion parameters were used for all systems.

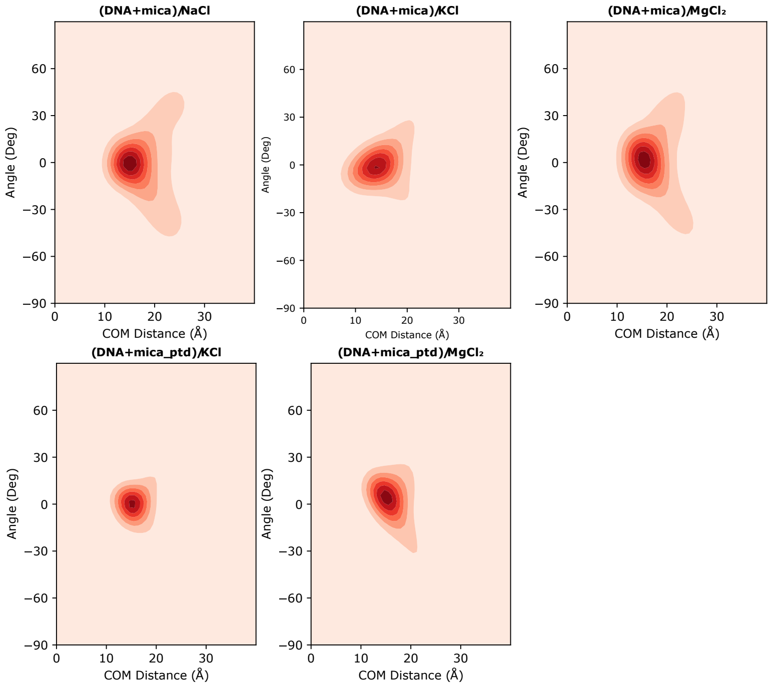


Figure S5: DNA angle distribution as a function of COM distance to DNA

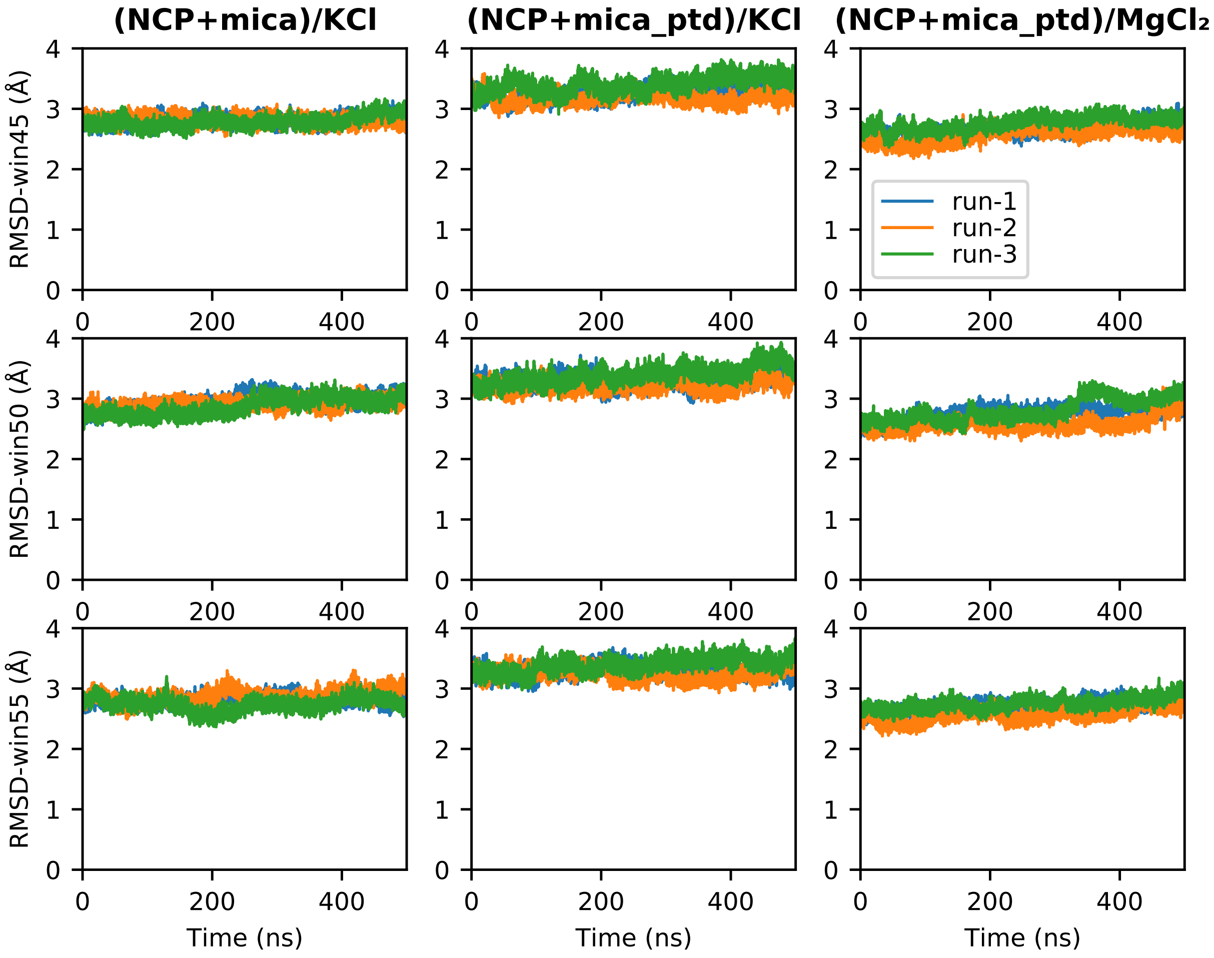


Figure S6: Core histone backbone RMSDs calculated for three systems at win45, win50 and win55.

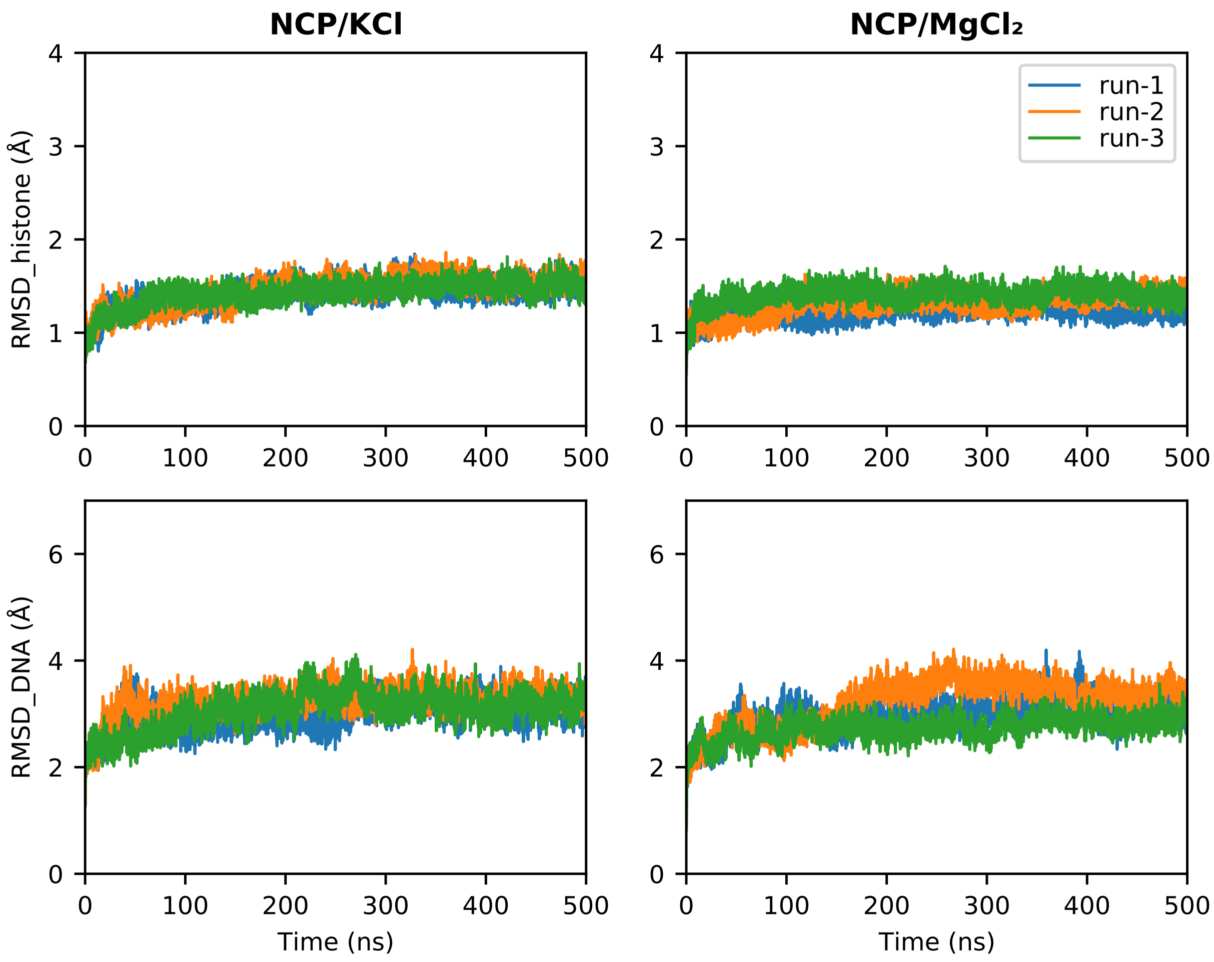


Figure S7: Histone core (Top) and DNA backbone (Bottom) RMSDs calculated for nucleosome in solution. Note the difference in Y-axis scale for histone and DNA RMSDs.

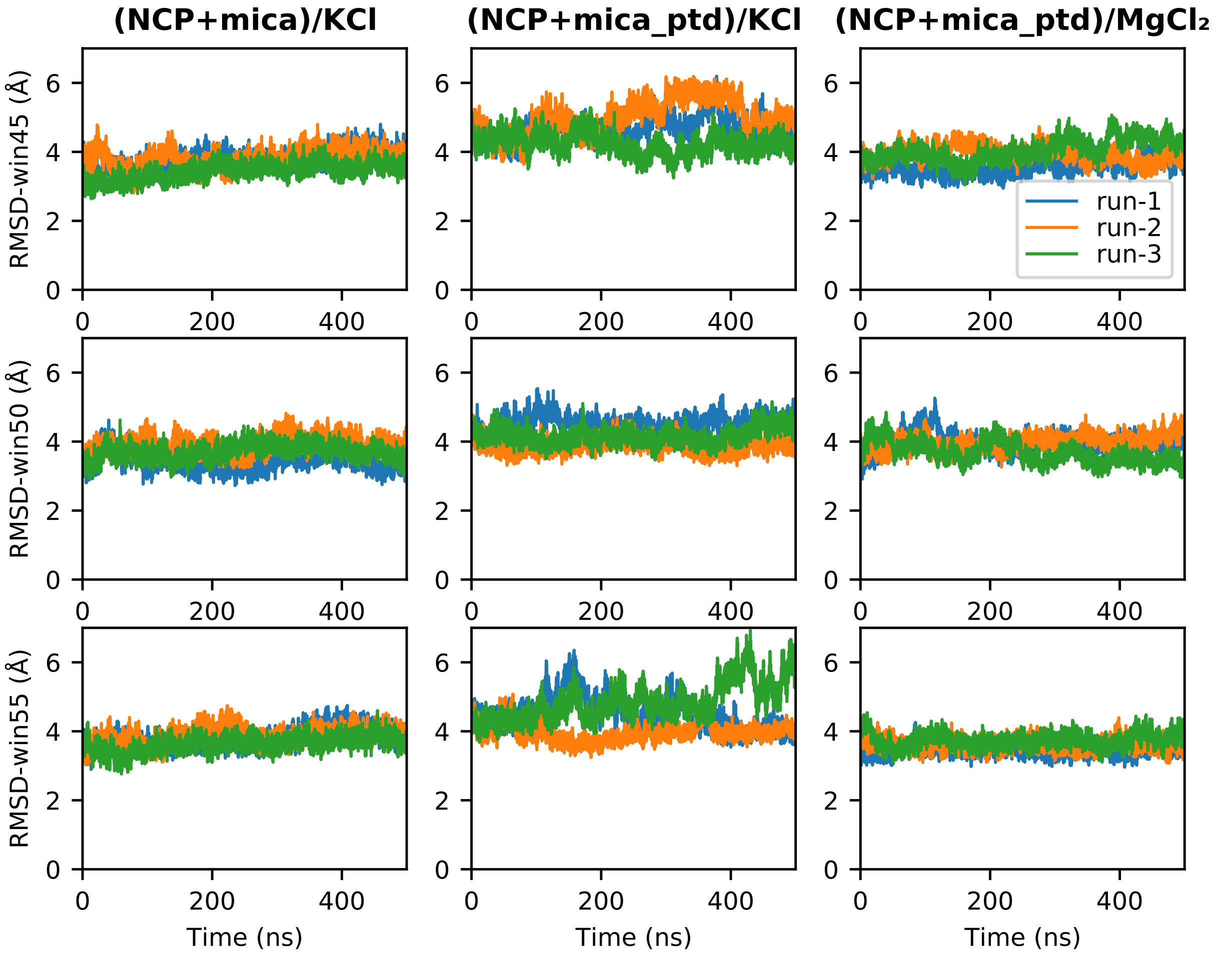


Figure S8: DNA backbone RMSDs calculated for nucleosome+mica systems at win45, win50 and win55.

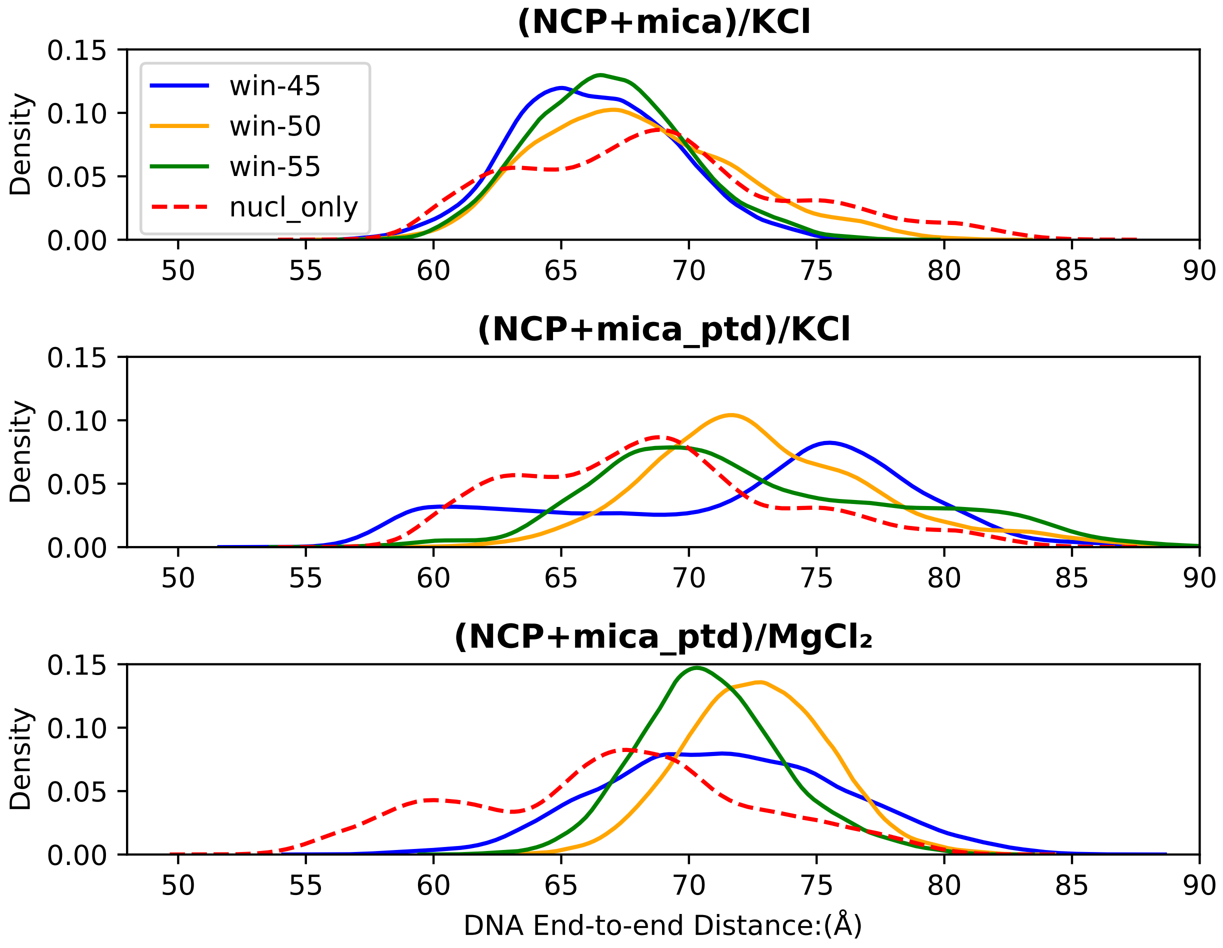


Figure S9: DNA end to end distance for each windows compared to the DNA end to end to end distance when nucleosome in solution without mica surface (red dashed line).

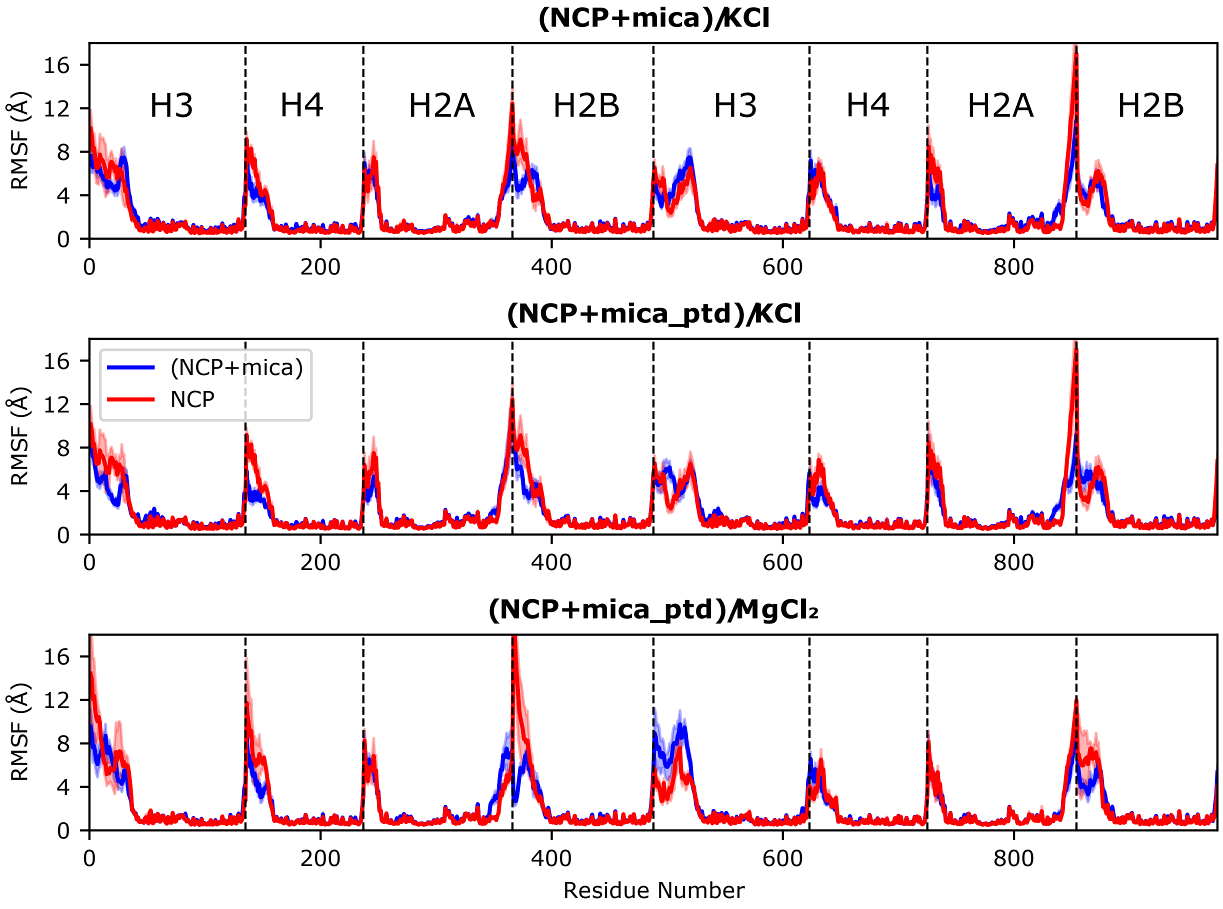


Figure S10: Histone residue RMSFs calculated for three nucleosome+mica systems are shown in blue and the nucleosome in corresponding solution (KCl or $MgCl_2$) are shown in red.

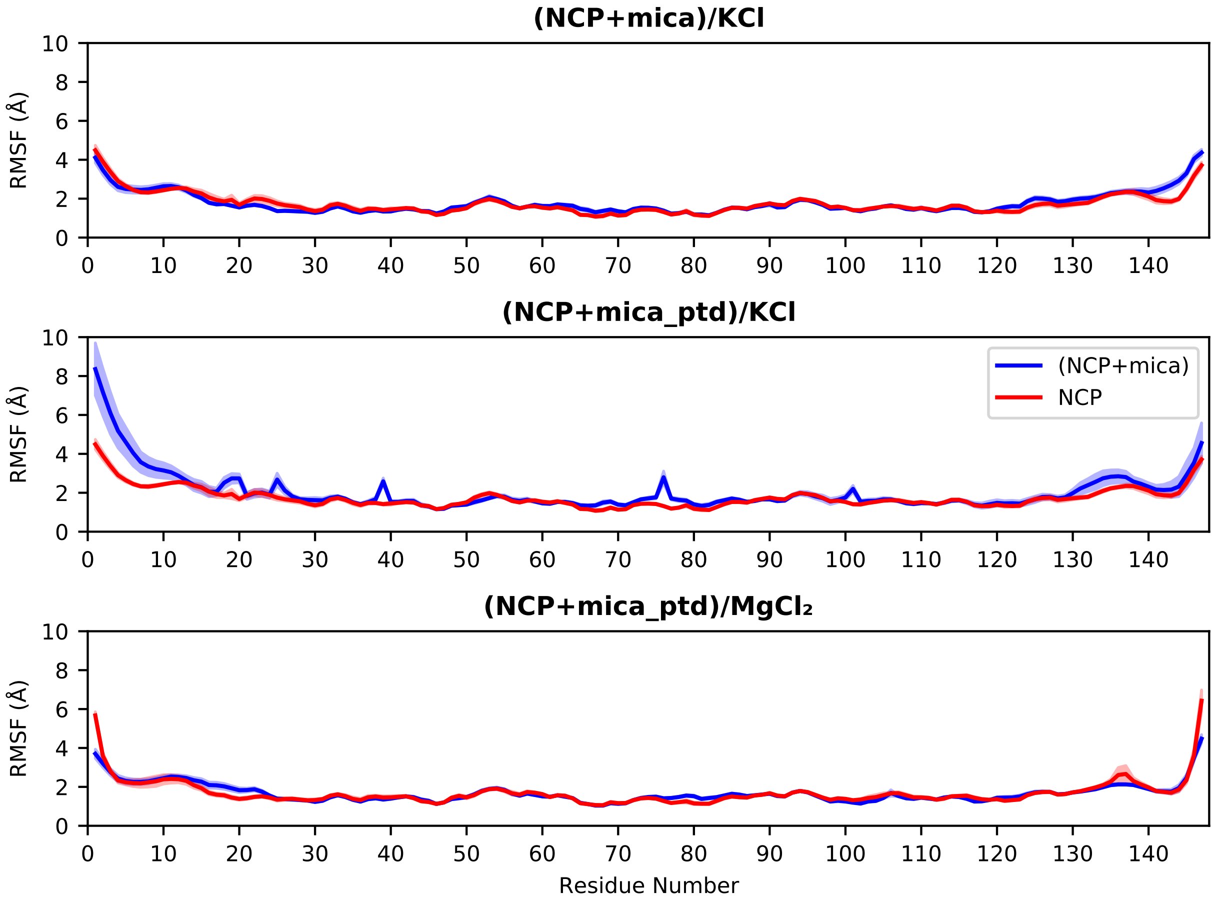


Figure S11: DNA base pair RMSFs calculated for three nucleosome + mica systems are shown in blue and for the nucleosome in corresponding solution (KCl or $MgCl_2$) are shown in red.

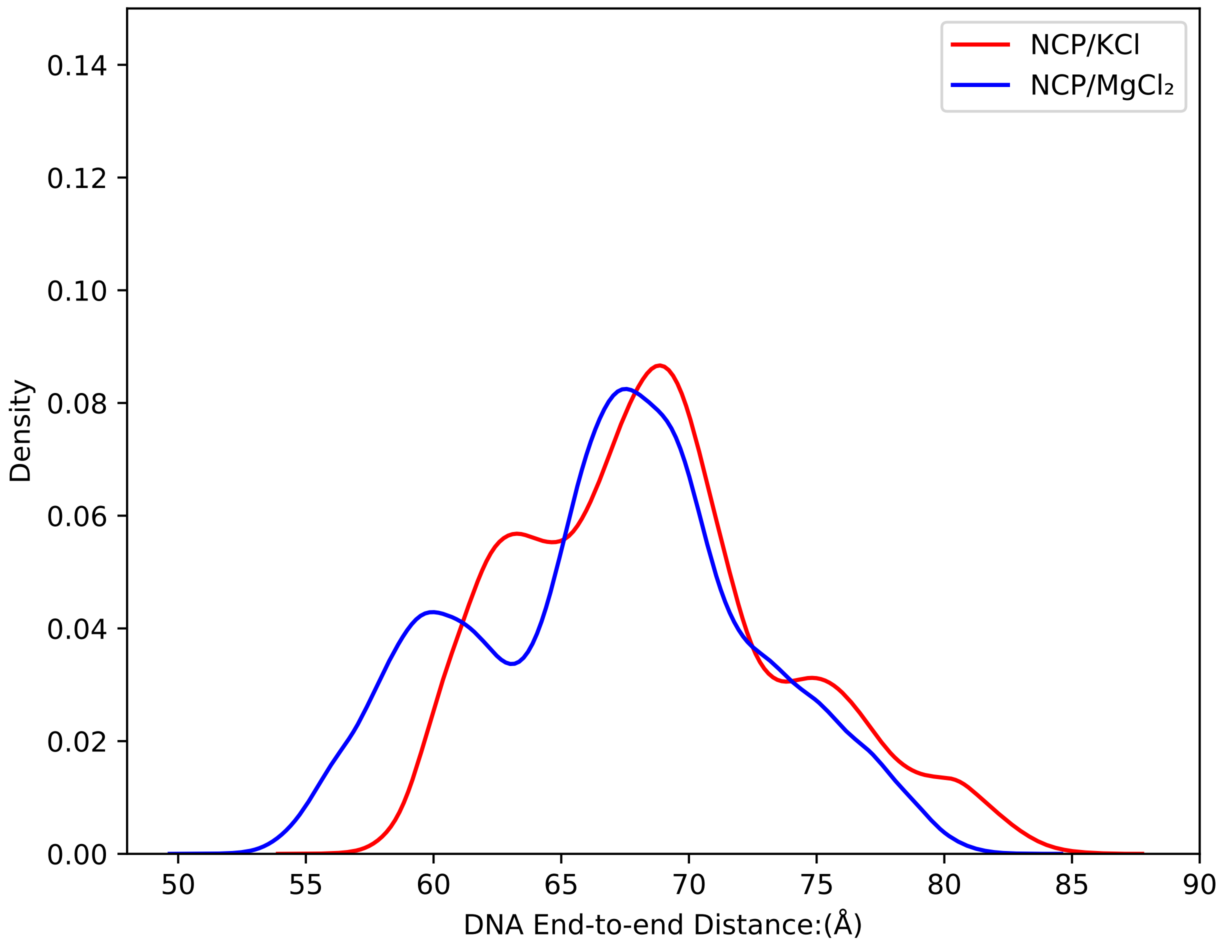


Figure S12: Comparison of DNA end to end distance when nucleosome in solution without mica surface.
